# Systems analysis of gut microbiome influence on metabolic disease in HIV and high-risk populations

**DOI:** 10.1101/2021.03.12.435118

**Authors:** Abigail J.S. Armstrong, Kevin Quinn, Jennifer Fouquier, Sam X. Li, Jennifer M. Schneider, Nichole M. Nusbacher, Katrina A. Doenges, Suzanne Fiorillo, Tyson J. Marden, Janine Higgins, Nichole Reisdorph, Thomas B. Campbell, Brent E. Palmer, Catherine A. Lozupone

## Abstract

Poor metabolic health, characterized by insulin resistance and dyslipidemia, is higher in people living with HIV and has been linked with inflammation, anti-retroviral therapy (ART) drugs, and ART-associated lipodystrophy (LD). Metabolic disease is associated with gut microbiome composition outside the context of HIV but has not been deeply explored in HIV infection nor in high-risk men who have sex with men (HR-MSM), who have a highly altered gut microbiome composition. Furthermore, the contribution of increased bacterial translocation and associated systemic inflammation that has been described in HIV-positive and HR-MSM individuals has not been explored. We used a multi-omic approach to explore relationships between impaired metabolic health, defined using fasting blood markers, gut microbes, immune phenotypes and diet. Our cohort included ART-treated HIV positive MSM with and without LD, untreated HIV positive MSM, and HR-MSM. For HIV positive MSM on ART, we further explored associations with the plasma metabolome. We found that elevated plasma lipopolysaccharide binding protein (LBP) was the most important predictor of impaired metabolic health and network analysis showed that LBP formed a hub joining correlated microbial and immune predictors of metabolic disease. Taken together, our results suggest the role of inflammatory processes linked with bacterial translocation and interaction with the gut microbiome in metabolic disease among HIV positive and negative MSM.

**Importance Statement:** The gut microbiome in people living with HIV (PLWH) is of interest as chronic infection often results in long term comorbidities. Metabolic disease is prevalent in PLWH even in well-controlled infection and has been linked with the gut microbiome in previous studies, but little attention has been given to PLWH. Furthermore, integrated analyses that consider gut microbiome together with diet, systemic immune activation, metabolites, and demographics have been lacking. In a systems-level analysis of predictors of metabolic disease in PLWH and men who are at high risk of acquiring HIV, we found that increased LBP, an inflammatory marker indicative of compromised intestinal barrier function, was associated with worse metabolic health. We also found impaired metabolic health associated with specific dietary components, gut microbes, and host and microbial metabolites. This work lays the framework for mechanistic studies aimed at targeting the microbiome to prevent or treat metabolic endotoxemia in HIV-infected individuals.

## Background

Poor metabolic health characterized by insulin resistance and dyslipidemia is frequent in people living with HIV (PLWH) (1–3) and has been linked with chronic inflammation (4–7) and several anti-retroviral therapy (ART) drugs (8). Metabolic disease is particularly prevalent in HIV-positive individuals with lipodystrophy (LD), a disease linked with early ART drugs that is manifested by lipoatrophy in the face, extremities, and buttocks with or without visceral fat accumulation. Delineating pathophysiological mechanisms of impaired metabolic health is crucial for tailoring strategies for prophylaxis and treatment to PLWH.

Metabolic disease has been linked with gut microbiome structure and function outside the context of HIV infection (9–13), but this relationship has not been explored deeply in PLWH. We and others have found an altered gut microbiome composition in both PLWH (14–16) and men who have sex with men at high-risk of contracting HIV (HR-MSM) (15, 17). Furthermore, we have demonstrated that the altered microbiome in HIV (14) and HR-MSM (14, 18) are pro-inflammatory *in vitro* and/or in gnotobiotic mice (14, 18). This is of interest as peripheral inflammatory signals have been implicated in both cardiovascular disease risk (7, 19) and insulin sensitivity (4, 5, 20–22) in PLWH. A recent study conducted in PLWH found that HIV-associated microbiome differences correlated with risk of metabolic syndrome, particularly in individuals with a history of severe immunodeficiency (23).

One mechanism by which the gut microbiome may impact MS in PLWH is through effects on intestinal barrier function. Impaired intestinal barrier, measured by increased bacterial products such as lipopolysaccharide (LPS; endotoxin) in blood, has been linked with MS and particular metabolic derangements (e.g., dyslipidemia and insulin resistance). This “metabolic endotoxemia” has been described in chronic kidney disease/ hemodialysis patients(24) and there are mixed data regarding a role in obesity(25–27). Murine studies have further supported that an altered gut microbiome and translocation of LPS could trigger insulin resistance, diabetes, and atherosclerosis(28–31). This is of interest for PLWH because bacterial translocation, indicated by higher levels of LPS or LPS-binding protein (LBP) in blood, is well known to occur in HIV infected populations, to be not completely ameliorated by antiretroviral therapy (ART)(32, 33), and to positively correlate with HIV-associated gut microbiome differences(23). Increased plasma LPS levels have also been observed in MSM and linked with recent sexual behavior(34).

We hypothesized that PLWH and HR-MSM with poor metabolic health would harbor a distinct gut microbial signature that was in turn also associated with elevated peripheral immune activation. We evaluated this relationship while considering other factors known to influence the microbiome, immunity and metabolic health. This analysis included typical diet; HIV, ART, and LD status; and other demographic characteristics such as age and body mass index (BMI). For HIV-positive individuals on ART with and without LD, we further explored associations with the plasma metabolome (Figure 1). Our results suggest a central role of inflammatory processes linked with bacterial translocation as measured by LBP, and co-correlated intestinal microbes, dietary and demographic attributes in metabolic disease risk.

**Figure 1.**
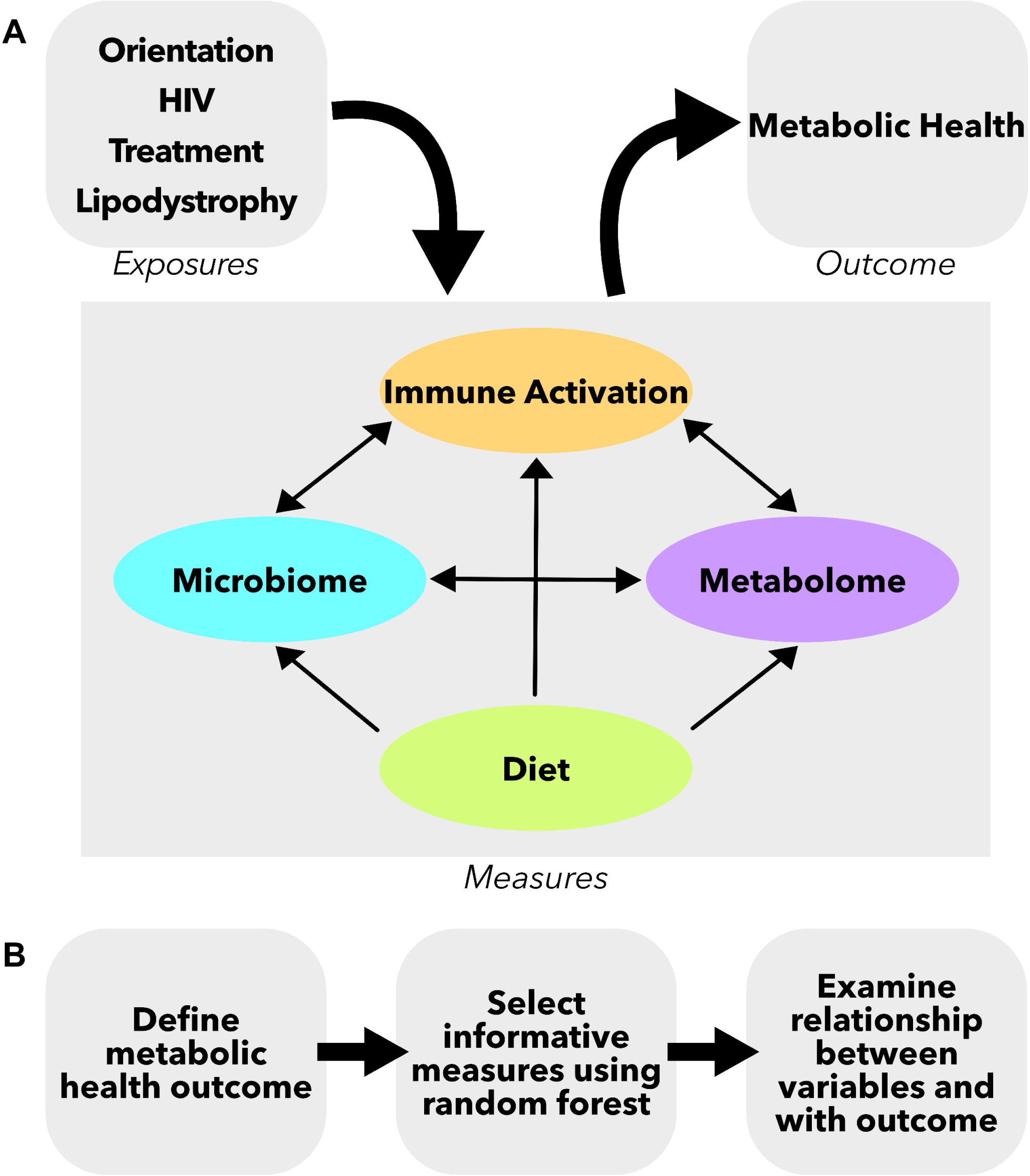
Study design schematic. **A.** Measures were collected from four compartments: gut microbiome, peripheral immune, diet questionnaire, and plasma metabolome. These separate compartments can all influence each other and can be influenced by other clinical and demographic characteristics such as HIV and treatment status. **B.** Analysis pipeline for the study. First, a metabolic health outcome was determined. Second, informative variables were selected using random forest analysis. Lastly, the relationships between these informative variables and the metabolic health outcome were examined.

## Results

### Study Population

This study examined a cohort of 113 men, including men who have sex with women (MSW; n=22, 19.5%) and MSM (n=91, 80.5%) (Table 1). Of the MSM, 32 were HIV-negative (35.2%), 14 were HIV-positive and not on ART (15.4%), and 45 were HIV-positive and on ART (49.4%). The HIV-positive, treated group included those with lipodystrophy (LD; n=25, 55.6%) and those without (n=20, 44.4%). The HIV-negative MSM participated in activities that put them at high risk of contracting HIV including: 1) a history of unprotected anal intercourse with one or more male or male-to-female transgender partners; 2) anal intercourse with two or more male or male-to-female transgender partners; or 3) being in a sexual relationship with a person who has been diagnosed with HIV (35). In order to focus on HIV-associated metabolic disease, obese individuals (BMI >30) were excluded. There was no significant difference in BMI between the cohorts (Kruskal-Wallis test, p = 0.085, Supplemental Table 1). Individuals in the HIV-positive, treated cohorts were significantly older than HIV-negative and HIV-positive, untreated MSM (Kruskal-Wallis test, p < 0.001). Age matching across all cohorts was not feasible in part because LD is associated with early-generation ART drugs and thus most common in older HIV-positive individuals and HR-MSM behavior and new HIV infections are predominantly in younger individuals. However, age is carefully considered in downstream analyses. All treated, HIV-positive individuals were on successful ART with suppressed viral loads (Table 1).

**Table 1.** Description of full study cohort.

|  | **HIV-negative**<br>**MSW** | **HIV-negative**<br>**MSM** | **HIV-positive**<br>**MSM,**<br>**untreated** | **HIV-positive**<br>**MSM,**<br>**treated** | **HIV-positive**<br>**MSM,**<br>**treated,**<br>**with LD** | **P** |
| --- | --- | --- | --- | --- | --- | --- |
| **n** | 22 | 32 | 14 | 20 | 25 |  |
| **Age**<br>**(years)** | 33<br>(27.3-38.5) | 34<br>(29.8-44.5) | 34<br>(26.5-40.3) | 46<br>(42.8-50.5) | 60<br>(54-64) | *** |
| **BMI (kg/m2)** | 25.2<br>(23.0-27.0) | 25.5<br>(20.2-28.0) | 21.4<br>(20.2-25.6) | 23.9<br>(22.6-26.2) | 25.8<br>(23.0-28.0) | NS |
| **CD4**<br>**cell count** | NA | NA | 538<br>(406-732) | 586<br>(420-878) | 659<br>(550-908) | NS |
| **CD4 nadir** | NA | NA | 496<br>(409-612) | 256<br>(118-416) | 152<br>(70-350) | *** |
| **Viral load** | NA | NA | 101,400<br>(20,300-292,514) | 20<br>(0-20) | 0<br>(0-20) | *** |
| **Cholesterol**<br>**drugs/statins**<br>**n (%)** | 2 (9.1%) | 3 (9.4%) | 1 (7.1%) | 4 (20%) | 14 (56%) | +++ |

### Metabolic Disease Score as a Marker for Metabolic Health

We measured seven common clinical markers of metabolic health from fasting blood: triglycerides, glucose, insulin, LDL, HDL, leptin, and adiponectin. Since these markers are often correlated with each other, we used principal component analysis (PCA) to define a single continuous measure of overall metabolic health as has been done previously (Figure 2A) (36, 37). Individuals with high values along the first principal component (PC1) generally had high triglycerides and low HDL, indicating dyslipidemia, and higher levels of fasting blood glucose and insulin, indicating insulin resistance (Figure 2A, Supplemental Figure 1). Values of PC1 were shifted to a minimum of one and log transformed to define the metabolic disease score, which ranged from 0 as healthy and 2.5 as impaired. We used regression analysis to determine how this score related to clinically defined cutoffs for normal levels of the markers (Supplemental Figure 1). For example, triglycerides positively correlated with metabolic disease score and almost all individuals with a score above 1.45 had triglyceride levels in the unhealthy range of greater than 200 mg/dL. Similar patterns and cutoffs were true for HDL, LDL, and glucose (Supplemental Figure 1). The intersect of the regression with these cutoffs were all averaged to a single number of 1.4. Individuals below the cutoff were categorized as metabolically normal and those above were categorized as metabolically impaired.

**Figure 2.**
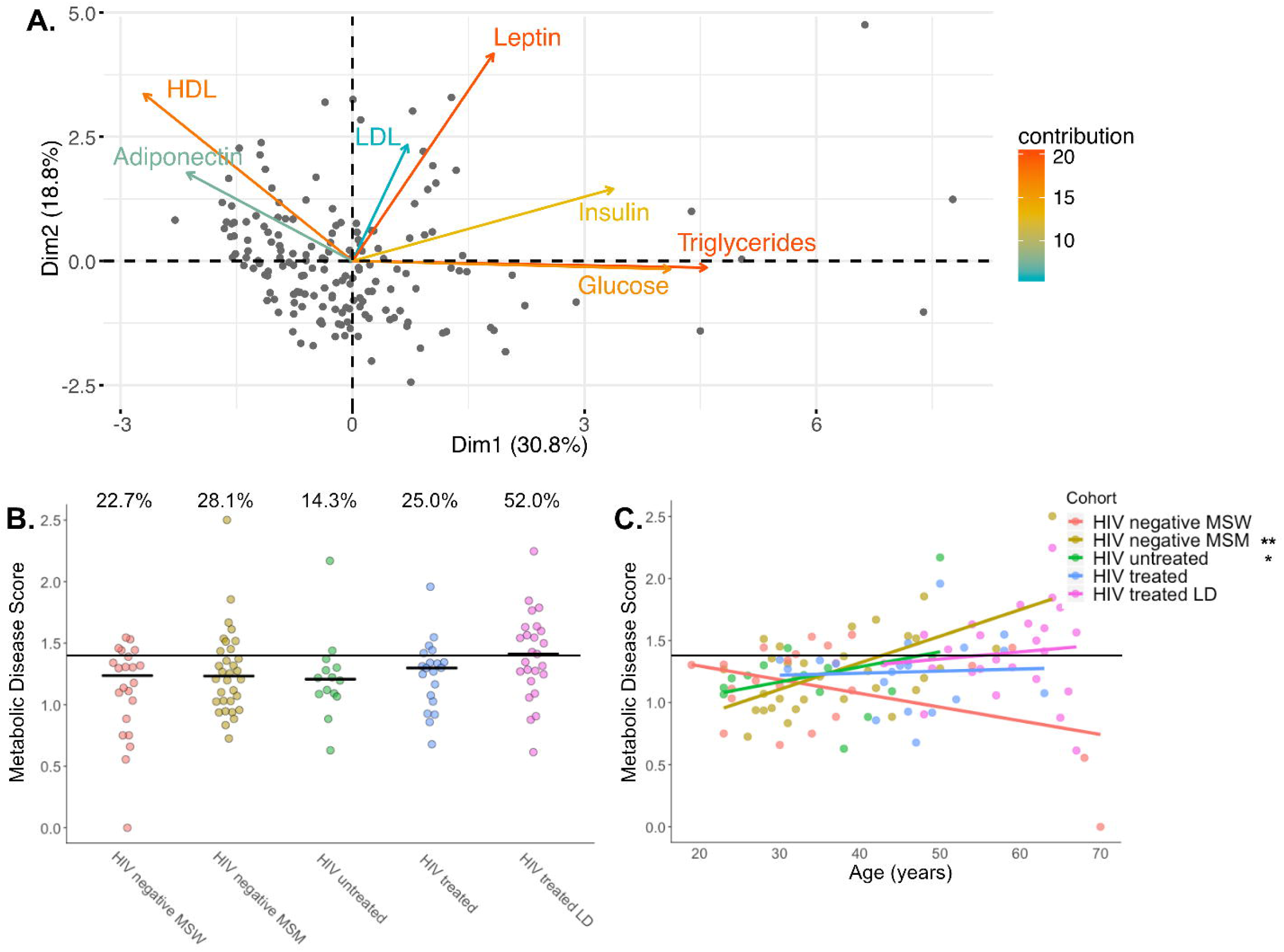
Calculation of the metabolic disease score. **A.** PCA of metabolic markers in fasting blood of 164 men and women: 113 participants described in this paper along with 51 individuals recruited at the same time and under the same exclusion criteria as study participants. Metabolic disease score is calculated as the PC1 coordinates shifted to a minimum of one and log transformed. **B.** Metabolic disease scores broken up by cohort. The percentages noted above the groups are the percent of individuals with a score above our metabolic impairment cutoff (Supplemental Figure 1). There is no significant difference between the proportions in each group (Fischer’s exact test, p = 0.11) or between mean ranks in each group (Kruskal-Wallis test p = 0.13). **C.** Relationships between metabolic disease score and age stratified by cohort. Statistical significance of slopes are indicated and were calculated with the linear model: score ∼ age + cohort + age*cohort. P-value annotations: ** < 0.01; * < 0.05.

When comparing the metabolic disease score across cohorts, we found that ART-treated, HIV-positive individuals with LD trended higher in both the average metabolic disease score and the proportion of individuals with scores in the metabolically impaired group but intergroup significance was lost after multiple test corrections (Figure 2B). Furthermore, because our HIV-positive, treated cohorts were significantly older than our HIV-negative MSM and HIV-positive, untreated MSM, we used a linear model to explore differences in the metabolic disease score across cohorts while accounting for age (Figure 2C). This score was positively associated with age only in HIV-negative MSM and HIV-positive, untreated MSM (Figure 2C; linear model; p < 0.001 and p = 0.036 respectively) and only HIV-negative MSM had significantly higher metabolic disease score compared to HIV-negative MSW when accounting for age (linear model; p < 0.001).

### Selection of Features that Predict the Metabolic Disease Score and Interactions Between Selected Features

We next explored the complex relationships of the gut microbiome, peripheral immune activation, and diet to the metabolic disease score and to each other using only data from the HIV positive and HR-MSM cohorts. We first selected features that were important predictors of the metabolic disease score using the VSURF (Variable Selection Using Random Forest) tool (38). VSURF is optimized for feature selection, returning all features that are highly predictive of the response variable, even when a smaller subset of highly predictive variables with redundant features removed could be just as accurate for prediction (38). We input the following features into the VSURF tool: 1) 130 microbial features. 99% identity Operational Taxonomic Units (OTUs) with highly co-correlated OTUs were binned into modules as described in the methods (detailed in Supplemental Table 2). Only OTUs present in >20% of samples were included. 2) 21 immune features that were measured in plasma using multiplex ELISA (detailed in Supplemental Table 2). These immune measures were selected based on a literature search for those previously shown to be altered in HIV infection and/or metabolic disease. 3) 21 clinical/demographic features such as age, BMI, HIV infection and treatment status, and typical gastrointestinal symptoms including constipation, diarrhea and bloating (detailed in Supplemental Table 2). 4) 29 dietary features that were collected using a food frequency questionnaire of typical dietary intake over the prior year as detailed in the methods. Highly co-correlated diet features were also binned into modules (detailed in Supplemental Table 2).

From the initial 201 measures, VSURF identified 69 important variables (4 clinical data measures, 6 diet measures, 14 immune measures, and 45 microbes) and a subset of ten highly predictive variables (Supplemental Table 3). These 69 features could predict the metabolic disease score using traditional random forest with an r^2^ of 31.05%, which is significantly better than a null model where the outcome was randomly permuted (p = 0.036, Supplemental Figure 2).

We found that 21 of the 69 selected variables were positively or negatively correlated with the metabolic disease score (Spearman rank correlation, FDR p < 0.1, Supplemental Table 3). Since random forest can detect non-linear relationships and/or features that are only important when also considering another feature, it is not surprising that all features were not correlated linearly with the metabolic disease score. All VSURF selected clinical measures were positively correlated with metabolic disease score and included age, BMI, lipodystrophy, and bloating (Supplemental Table 3). None of the six selected diet measures correlated with metabolic disease score (Supplemental Table 3). VSURF selected immune markers that were positively correlated with the metabolic disease score included LBP, intercellular adhesion molecule 1 (ICAM-1), interleukin (IL) 16, IL-12, and granulocyte-macrophage colony-stimulating factor (GM-CSF) (Supplemental Table 3). The feature with the highest random forest importance score was LBP.

Diet, the microbiome, and immune phenotypes can all influence each other and can also relate to clinical/demographic factors such as BMI and age (Figure 1). For this reason, we also investigated the relationship between the 69 important factors using pairwise Spearman rank correlation and network visualization (Figure 3, Supplemental Figure 3, Supplemental Table 3). The selected important microbes included many that were highly correlated with each other and with dietary, clinical/demographic, and inflammatory phenotypes (Supplemental Figure 3A, Supplemental Table 3). For example, a module of bacteria identified within the *Prevotella* genus and the *Paraprevotellaceae* family, negatively associated with metabolic disease score and positively associated with dietary fiber (Supplemental Figure 3B).

**Figure 3.**
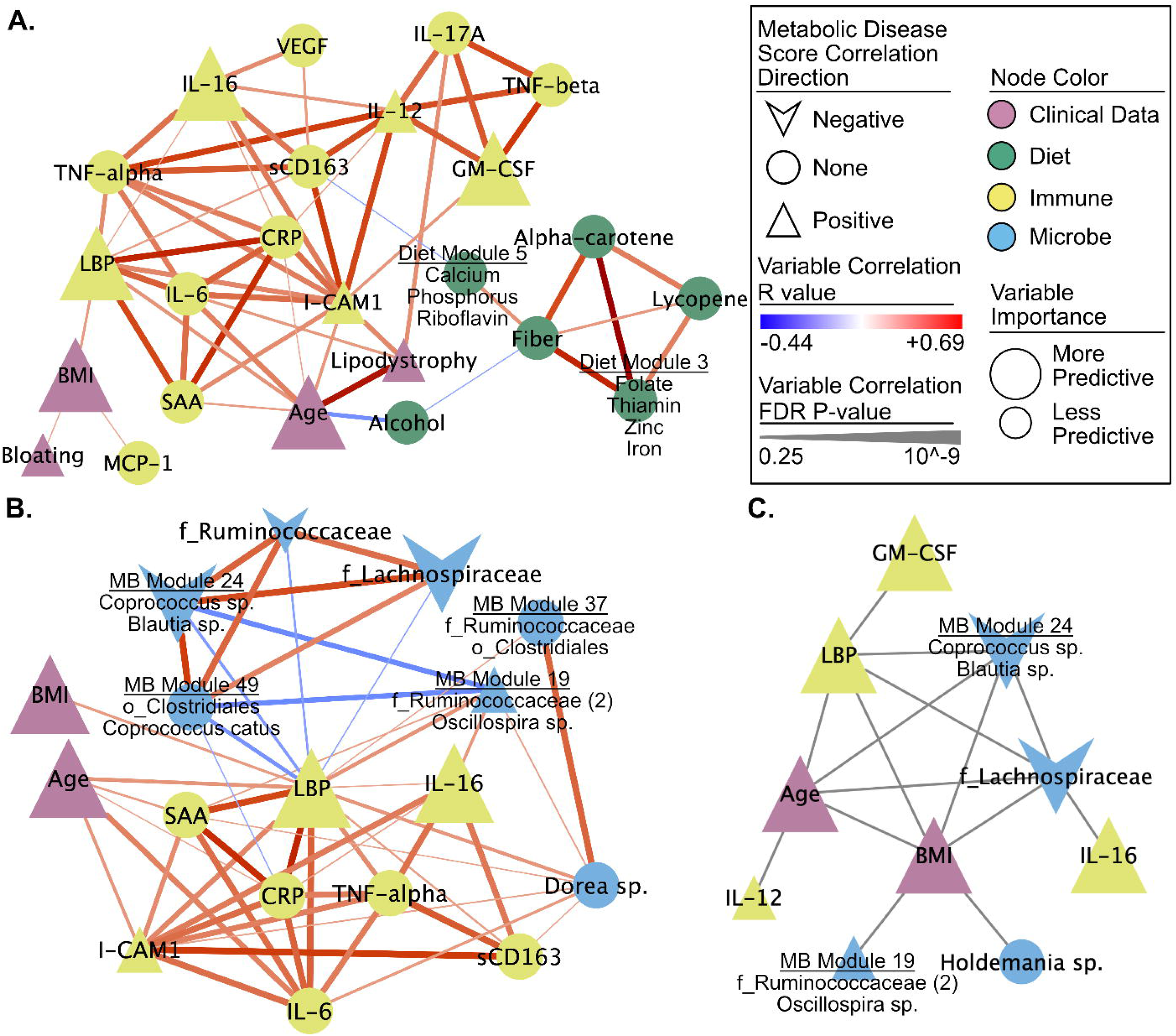
Networks of selected measures reveal several strong associations with metabolic disease score and between measures. Correlation sub-networks of **A.** all the non-microbe selected measures. **B.** the nearest neighbors of LBP. All Spearman rank correlations with an FDR p < 0.25 are shown. Subnetworks were pulled from a larger network of all VSURF selected measures (Supplemental Figure 3, Supplemental Table 2). **C.** Network of interactions between measures calculated using iRF. All edges represent an interaction (i.e., proximity in a decision tree) that occurred in 30% or more of the decision trees.

Because bacterial translocation is known to occur at increased levels in both HIV-positive individuals (39) and HR-MSM (34), we were specifically interested in investigating which of the selected features correlated with LBP. LBP was negatively correlated with several putative butyrate producing bacteria/bacterial modules such as OTUs in the genera *Coprococcus* (40, 41)(Figure 3B). LBP was also positively correlated with *Dorea* species (Figure 3B; Supplemental Table 3). In addition to correlations, we evaluated interactions between selected variables using the tool iRF (iterative Random Forest)(42). These interactions represent variables that are in adjacent nodes in a random forest tree in which the value of one influences the predictability of the other. This analysis also identified that LBP interacted with age, BMI, an OTU in the *Lachnospiraceae* family, and microbiome module 24 (*Coprococcus* sp. and *Blautia* sp.) in predictions of the metabolic disease score (Figure 3C; Supplemental Table 3).

Because of the recognized role of inflammation in an increased risk of age-associated non-AIDS-related morbidity and mortality in PLWH(43–46), we were also specifically interested in investigating which of the selected features correlated with age. Age had the strongest positive association with lipodystrophy and also positively correlated with several inflammatory markers including LBP, CRP, IL-6, ICAM-1 and SAA (Supplemental Figure 3C, Supplemental Table 2). Age was also correlated with several different gut microbes including a negative correlation with OTUs assigned to *Bifidobacterium adolescentis*, *Eubacterium dolichum, Coprobacillus sp.*, and *Oscillospira sp.* (Supplemental Figure 3C).

Random Forest analyses do not allow for gaps in data, and, as a result, HIV-specific variables were omitted from these analyses. We thus performed Spearman’s rank correlations to investigate a relationship between variables only measured in our HIV positive cohorts and the metabolic disease score and found no correlation with CD4 nadir (p=0.28) and CD4+ T-cell count (p=0.076). We also found no significant correlation between viral load and metabolic disease score among individuals with untreated HIV infection (p = 0.68).

Lastly, because many gut microbes were associated with metabolic disease score, we also investigated associations with overall microbiome composition measures. Microbiome evenness, as measured by Pielou’s Evenness and Shannon Index were both weakly negatively correlated (Spearman Rank Correlation, rho=-0.09; p = 0.047 and rho=-0.09; p = 0.045 respectively) and there was no significant relationship in measures of microbiome richness (Faith’s PD and Observed Features). Additionally, there was no significant relationship between weighted and unweighted UniFrac distances and metabolic disease score differences between individuals (Mantel Test, p = 0.7 and p = 0.4 respectively).

### The Relationship Between The Plasma Metabolome and the Metabolic Disease Score in ART-treated HIV-positive Individuals With and Without LD

To pursue a further mechanistic understanding of how the gut microbiome may influence the metabolic disease score in PLWH, we performed untargeted metabolomics (LC/MS) on plasma from our cohort of ART-treated, HIV-positive individuals with and without LD (n=44). Metabolite identities were then validated using untargeted MS/MS. We used two approaches to determine which plasma metabolites were either directly produced or indirectly influenced by the gut microbiome. First, metabolites were run through the computational tool AMON (47), which uses the KEGG database (48) and inferred metagenomes (calculated using PICRUSt2 (49)) to determine which could have been produced by the microbiome. Second, LC/MS was run on plasma from both germ free (GF) and humanized mice to determine metabolites that had significantly altered levels upon colonization with human microbiomes. Specifically, GF mice were gavaged using fecal samples from eight men from the study cohort (humanized mice) while two mice were gavaged using PBS as control (Supplemental Table 4). Plasma was collected before and after gavage. All mice were fed a high-fat western diet.

We found that 820 metabolites were different in abundance between GF and humanized mice after multiple test corrections (Student’s t-test, FDR p < 0.05), 493 of which were also present in the human plasma samples (Figure 4). From the full set of 5,332 metabolites identified in the human plasma, 416 could be annotated with KEGG IDs. These were further analyzed using AMON. 146 microbiome-associated metabolites were identified that are putatively produced by the gut microbiome; however, many of these could also be produced by the host. Twenty-six of the 146 microbiome-associated metabolites identified by AMON also differed in colonized versus GF mice (Figure 4, Supplemental Table 5).

**Figure 4.**
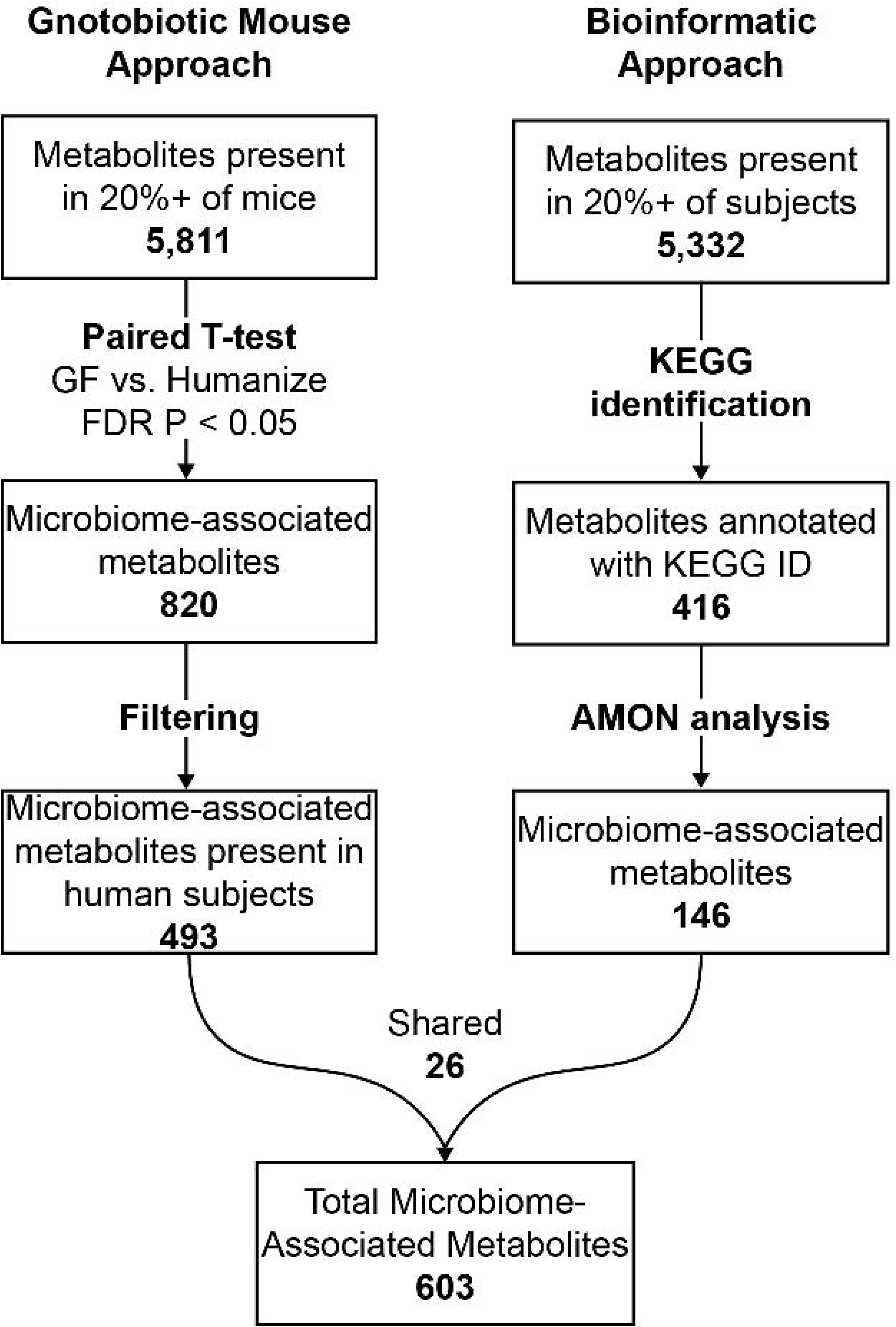
Microbiome-associated metabolite workflow. Two-pronged approach of identifying microbiome-associated metabolites. Numbers in bold are counts of metabolites.

Of the 5,332 total measured metabolites in the human samples, 150 correlated with the metabolic disease score (Spearman rank correlation, FDR p < 0.05; Supplemental Table 5). The correlated compounds were enriched in a number of different metabolic pathways with both the Phospholipid and the Glycerolipid pathways of the Small Molecule Pathway Database (SMDPH)(50) highly enriched (Supplemental Table 5). Consistent with the metabolic disease score being defined in part by dyslipidemia, 17 of the significant compounds were annotated as triglycerides. Seven of the significant compounds were associated with the microbiome as determined by the process outlined above. We confirmed the identity of 5 of the 7 of these with MS/MS (Supplemental Table 5). Of these seven microbiome-associated metabolites, two could exclusively be explained by direct production by the microbiome. Specifically, dehydroalanine, identified as a microbial product by AMON, negatively correlated with the metabolic disease score and bacteriohopane-32,33,34,35-tetrol positively correlated with the metabolic disease score (Supplemental Table 5). Two additional microbiome-associated compounds were triglycerides (TG(54:6) and TG (16:0/18:2.20:4)) that were positively correlated with metabolic disease score and elevated in humanized mice compared to GF mice. Another of these metabolites, 1-Linleoyl-2-oleoyl-rac-glycerol, is a 1,2-diglyceride in the triglyceride biosynthesis pathway. Finally, phosphatidylcholine (PC(17:0/18:2)) and phosphatidylethanolamine (PE(20:3/18:0)) compounds were identified as microbiome associated and positively and negatively correlated with the metabolic disease score respectively (51).

Since age was significantly associated with metabolic disease score in the full cohort, and thus could present a potential confounding factor in the metabolite relationships, we constructed a model that included age using a rank-based estimation for linear models using the R package Rfit (52). Because of the statistical differences between linear regression and spearman correlation, we looked at Rfit with and without age. Both analyses identified 60 metabolites significant after FDR correction (Supplemental Table 5) and 45 of the 60 metabolites were identified with both Rfit models, indicating that age was not very influential on the results (Supplemental Figure 4).

## Discussion

In this study, we identified gut microbes, dietary components, demographic and immune measures that predicted impaired metabolic health in a cohort of MSM with and without HIV, ART, and LD. Notably, we identified a strong relationship with circulating LBP which in turn correlated with other markers of systemic inflammation, a loss of beneficial microbes such as butyrate-producing bacteria, and higher BMI, indicating that diverse modifiable factors may influence LPS/inflammation driven metabolic disease in this population.

There was a positive association between impaired metabolic health and age, as has been reported previously for both HIV (53) and non-HIV populations (54), but linear modeling suggested that this relationship was driven by an association in HIV-negative MSM and HIV-positive untreated MSM in our study, revealing a possibly larger effect size than in our other cohorts. Also, when controlling for age, HIV-negative MSM had the highest metabolic disease score, even compared to HIV positive individuals on ART with LD, a population that has previously been reported to have higher incidence (55). This result is intriguing given prior research linking increased levels of LPS in blood with high-risk behavior in MSM (34). Larger cohorts and more detailed behavior information are required, however, to make any definitive claims on impaired metabolic health in aging in HR-MSM.

Our finding that age was a significant predictor of impaired metabolic health in this cohort is consistent with previous studies showing an increased risk of age-associated non-AIDS-related morbidity and mortality in PLWH (44–46, 56). Consistent with the previously reported role of inflammation in this relationship, age was positively correlated with 5 of the 14 immune measures that VSURF selected as important predictors of the metabolic disease score, including LBP, CRP, IL-6, ICAM-1 and SAA. Consistent with age being associated with changes in the microbiome previously (57–59), we also found age to be correlated with several different predictive microbes (Supplemental Figure 3C, Supplemental Table 2). These included a negative association with *B. adolescentis*, whose loss has previously been mechanistically linked with health deficits that occur with aging(60–63). *B. adolescentis* has been shown to prevent immunosenescence when fed probiotically to aged rats (62), and to have beneficial effects on barrier function in a murine model(64).

Consistent with prior studies that have associated high BMI with dyslipidemia, insulin resistance, and/or metabolic syndrome (65–67), BMI was a positive predictor of impaired metabolic health in our cohort even though our study excluded obese individuals, but did include overweight. This suggests the importance of weight management even among overweight, non-obese individuals as a strategy for reducing metabolic health impairment in this population.

We did not find a positive association between ART status and the metabolic disease score, but this may be because study participants were on a wide variety of drug combinations with the potential to have varied/contrasting effects. For instance, both integrase stand transfer inhibitors (ISTI) (68) and regimens including the nucleoside reverse transcriptase inhibitor (NRTI) tenofovir have been shown to increase risk of weight gain (69). Conversely, the CCR5 antagonist, maraviroc, may confer a benefit to cardiovascular function and body weight maintenance and evidence in mice suggests that these beneficial effects may be linked to gut microbiome composition changes (20, 70). Thus, future studies will be required to understand factors important in particular drug contexts. Other HIV-specific measures such as CD4 nadir, viral load and CD4+ T cell count also did not have a significant association with metabolic disease score. Low CD4+ T cell count (71) and CD4/CD8 T cell ratio (72) has previously been linked with poor metabolic health in individuals on ART. Furthermore, low CD4 nadir has been linked with gut microbiome dysbiosis in HIV-infected individuals (71) and an association between HIV-associated gut microbiome dysbiosis and metabolic syndrome was significantly stronger in individuals with past severe immunodeficiency compared to those without (23). Our negative results may be a result of small sample size and also a lack of individuals included in the study with severely compromised CD4+ T cell counts (Table 1).

Several dietary components that we identified have been previously associated with metabolic health, including dietary carotenoid, lycopene, and fiber (73–77). Fiber’s benefit in glucose response has been linked with the activity of *Prevotella copri*. Specifically, individuals who had improved glucose response after 3 days of high-fiber consumption had a greater increase in *P. copri*, and these beneficial effects were confirmed in a mouse model (76). However, another study found that *P. copri* actually promoted poor glucose response in the context of a western diet low in fiber through the production of branched chain amino acids (BCAAs)(11). Interactions between *Prevotella*, dietary fiber, and metabolic health are of particular interest in this cohort since HIV positive and negative MSM have much higher *Prevotella*, including *P. copri*, than non-MSM (15, 17). Additionally, our prior study using *in vitro* stimulations of human immune cells with fecal bacteria of HIV positive and negative MSM indicated that the Prevotella-rich microbiomes of MSM could drive systemic inflammation (14). Interestingly, in our data, a module of three OTUs, two in the genus *Prevotella* and one in the family *Paraprevotellaceae,* negatively associated with metabolic disease score and positively associated with dietary fiber (Supplemental Figure 3B), supporting a relationship between *Prevotella* and dietary fiber in improving metabolic health, and not supporting deleterious effects. Further work will be needed to decompose the complex relationship between dietary fiber, particular *Prevotella* strains, and metabolic health in HIV positive and negative MSM with unique Prevotella-dominated communities.

LBP was the most important feature in the random forest analysis and also a highly interactive measure in the iRF analysis. LBP binds to both microbial LPS and lipoteichoic acid (LTA) (78) and the presence of elevated LBP in blood is indicative of increased intestinal barrier permeability (79). LBP was correlated with age and BMI, a relationship that was previously observed in a cohort of HIV-negative men of African ancestry with this trio being further associated with adiposity and pre-diabetes (80). LBP levels were also correlated with other inflammatory markers that have been linked with worse metabolic health suggesting a role as a central mediator of metabolic-disease associated immune phenotypes. These included 1) I-CAM 1, whose expression in adipose tissue has been associated with diet-induced obesity in mice (81) and metabolic syndrome in humans (82) 2) IL-6, a pro-inflammatory cytokine that has been shown to play a direct role in insulin resistance (83), and 3) SAA, which is regulated in part by IL-6 and plays a role in cholesterol metabolism (82); SSA3 specifically has been shown to be produced in response to gut bacteria in obesity in mice (84). We observed a positive association between the metabolic disease score and frequency of abdominal bloating (Supplemental Table 3, Supplemental Figure 3A), further supporting a role of intestinal dysfunction in this population. Taken together these associations suggest that inflammation originating from an impaired intestinal barrier is promoting worse metabolic health.

An importance of microbiota-driven intestinal barrier dysfunction in HIV-associated metabolic syndrome was suggested in a recent study of Gelpi *et al.,* which found that metabolic syndrome in HIV-infected individuals was correlated with an HIV-associated gut microbiota (23). This microbiota was characterized in part by a decrease in butyrate-producing bacteria, including *Coprococcus* and *Butyrivibrio*. Consistent with this finding, we found that multiple *Coprococcus* OTUs were selected as important predictors of the metabolic disease score and that they also negatively correlated with LBP. Butyrate is well known to have beneficial effects on intestinal barrier function (85–94), and low levels of intestinal butyrate producers has been previously associated with microbial translocation and immune activation in PLWH (95). The prior study of Gelpi *et al.* (23) also found increased levels of bacteria in the *Desulfovibrionaceae* family to be linked with HIV-associated metabolic syndrome; These bacteria produce hydrogen sulfide, a compound that can compromise the intestinal mucus layer by reducing disulfide bonds (96). Further supporting that microbes that may compromise the mucus layer can impact barrier function in this context, we found *Dorea sp*, which encodes the genes required to utilize the canonical sialic acid Neu5Ac as a carbon source (97), to be an important predictor of the metabolic disease score and to positively correlate with LBP. Hypo-sialylated intestinal glycans have previously been linked with higher abundance of glycan-degrading species and higher levels of microbial translocation in HIV-infected individuals (98). Taken together, these results suggest that impaired metabolic health in HIV positive and negative MSM may be influenced by impaired intestinal barrier linked with both protective and detrimental metabolic activity of intestinal bacteria.

In our metabolomic analysis, we identified 150 metabolites in blood that correlated with the metabolic disease score. To identify compounds whose prevalence may be related to the gut microbiome we used two complimentary approaches. First, we used the bioinformatics tool AMON (47), which allows us to specifically evaluate which compounds could have been directly produced by the gut microbiome but is limited by a lack of KEGG annotations for many compounds. Second, we measured which compounds changed in relative abundance in GF versus mice colonized with feces from our study cohort, which can identify microbial influence in unannotated compounds but cannot differentiate between direct production/consumption by microbes versus indirect influence. These results will also be influenced by physiological differences between mice and humans and incomplete colonization. Although these weaknesses may have led us to underestimate which of the 150 metabolic disease associated compounds may have been related to the microbiome, it still identified compounds that supported a mechanistic link between gut microbes, metabolites, and metabolic disease in HIV-infected individuals on ART.

First, we found a negative correlation between the microbially-produced non-canonical amino acid, dehydroalanine, and metabolic disease score. Dehydroalanine is a component of lantibiotics that are active against Gram-positive bacteria. Second, we found a positive correlation with bacteriohopane-32,33,34,35-tetrol. This compound is a lipoxygenase inhibitor that prevents the formation of hydroxyicosatetraenoic acid and various leukotrienes from arachidonic acid (99), which have been linked with the development of cardiovascular disease and metabolic syndrome (100). This association of a potentially protective metabolite with an increase in metabolic impairment seems counterintuitive; however, it may be indicative of larger systemic changes in arachidonic acid metabolism. Third, we identified a PC and a PE associated with both the microbiome and metabolic disease score. Changes in PCs and/or PEs have been previously implicated in atherosclerosis, insulin resistance and obesity (51). AMON analysis indicated that both PCs and PEs can be synthesized by intestinal bacteria; however, these compounds can also be synthesized in the host and may be found in the diet. In our analysis, however, PE(18:1/20:1) levels were higher in colonized compared to GF mice indicating that intestinal bacteria do influence overall levels despite diverse potential sources. Finally, we observed increased levels of several plasma triglycerides in the humanized compared to GF mice, including two plasma triglycerides that were significantly associated with metabolic disease score. This confirms the influence of the gut microbiome on host plasma triglycerides (101–103). However, we did not find any strong associations between these triglycerides and specific microbes within our dataset, indicating a potential need for studies conducted in larger cohorts or with shotgun metagenomics to look for functional correlates.

In conclusion, we observed a relationship between diet, gut microbiome, plasma metabolome, and peripheral immune markers of inflammation and metabolic disease in HIV positive and HR-MSM. Our results suggest a central role of inflammatory processes linked with bacterial translocation and interaction with the gut microbiome, age and BMI in metabolic disease among HIV positive and negative MSM. Our results also suggest contributions of low fiber, key vitamins, and microbially produced metabolites. These results illuminate potential microbiome-targeted therapies and personalized diet recommendation given an interacting set of gut microbes and other host factors. Understanding these relationships further may provide novel treatments to improve the metabolic disease and inflammatory outcomes of MSM living with HIV.

## Methods

### Subject Recruitment

Participants were residents of the Denver, Colorado metropolitan area and the study was conducted at the Clinical Translational Research Center of the University of Colorado Hospital. The study was reviewed and approved by the Colorado Multiple Institutional Review Board and informed consent was obtained from all participants. For detailed criteria on recruitment of our five cohorts (HIV negative MSW; HIV negative MSM; HIV positive, ART naïve MSM; HIV positive ART-treated MSM with LD; and HIV positive ART-treated MSM without LD) see supplemental methods.

Feces, a fasting blood sample, and clinical surveys were collected from participants in order to obtain analytes for the study design outlined in Figure 3 (Supplemental Table 2). Additional information about relevant clinical measures such as probiotic use were collected via a questionnaire and study participants also filled out information on typical frequency of high-risk sexual practices and on typical levels of gastrointestinal issues such as bloating, constipation, nausea and diarrhea.

### Diet Data FFQ Collection

Typical dietary consumption over the prior year was collected using the Diet History Questionnaire II (104). Diet composition was processed using the Diet*Calc software and the dhq2.database.092914 database (105). All reported values are based on USDA nutrition guidelines. Reported dietary levels were normalized per 1000 kcal. To reduce the number of comparisons within the diet survey data, we binned highly co-correlating groups of measures within the data types into modules (Supplemental Table 2). These modules were defined using the tool, SCNIC (106).

### Immune Data Collection

Whole blood was collected in sodium heparin vacutainers and centrifuged at 1700rpm for 10 minutes for plasma collection. Plasma was aliquoted into 1mL microcentrifuge tubes and stored at −80°C. For ELISA preparation, plasma was thawed, kept cold, and centrifuged at 2000xg for 20 minutes before ELISA plating. Markers for sCD14, sCD163, and FABP-2 were measured from plasma using standard ELISA kits from R&D Systems (DC140, DC1630 &DFBP20). Positive testing controls for each ELISA kit were also included (R&D Systems QC20, QC61, & QC213). LBP was measured by standard ELISA using Hycult Biotech kit HK315-02. Markers for IL-6, IL-10, TNF-α, MCP-1, and IL-22 were measured using Meso Scale Discovery’s U-PLEX Biomarker Group 1 multiplex kit K15067L-1. Markers for SAA, VCAM-1, ICAM-1, and CRP were measured using Meso Scale Discovery’s V-Plex Plus Vascular Injury Panel 2 multiplex kit K15198G-1. Vascular Injury Control Pack 1 C4198-1 was utilized as a positive control for this assay. Markers for GM-CSF, IL-7, IL-12/23p40, IL-15, IL-16, IL-17A, TNF-β, and VEGF were measured using Meso Scale Discovery’s V-Plex Plus Cytokine Panel 1 multiplex kit K151A0H-1. Cytokine Panel 1 Control Pack C4050-1 was utilized as a positive control for this assay. Plasma samples were diluted per manufacturer’s recommendation for all assays. Standard ELISA kit plates were measured using a Vmax® Kinetic Microplate Reader with Softmax® Pro Software from Molecular Devices LLC. Multiplex ELISA kits from Meso Scale Discovery were measured using the QuickPlex SQ 120 with Discovery Workbench 4.0 software.

### Gnotobiotic Mouse Protocols

Germ-free C57/BL6 mice were purchased from Taconic and bred and maintained in flexible film isolator bubbles, fed with standard mouse chow. Three days before they were gavaged, male mice between 5-7 weeks of age were switched to a western high-fat diet and were fed this diet for the remainder of the experiment. Diets were all obtained from Envigo (Indiana): Standard chow-Teklad global soy protein-free extruded (item 2920X - https://www.envigo.com/resources/data-sheets/2020x-datasheet-0915.pdf), Western Diet – New Total Western Diet (item TD.110919). See Supplemental Table 4 for detailed diet composition. Mice were gavaged with 200 μL of fecal solutions prepared from 1.5 g of donor feces mixed in 3 mL of anaerobic PBS (18). Mice were housed individually following gavage for three weeks in a Tecniplast iso-positive caging system, with each cage having HEPA filters and positive pressurization for bioexclusion. Feces were collected from mice at day 21 for 16S rRNA gene sequencing. Mice were euthanized at 21 days post gavage using isoflurane overdose and all efforts were made to minimize suffering. Blood from euthanized animals was collected using cardiac puncture and cells were pelleted in K2-EDTA tubes; plasma was then aliquoted and stored at −80° C.

### Metabolomics Methods

#### Plasma Sample Preparation

A modified liquid-liquid extraction protocol was used to extract hydrophobic and hydrophilic compounds from the plasma samples (107). Briefly, 50 µL of plasma spiked with internal standards underwent a protein crash with 250 µL ice cold methanol. 750 µL methyl tert-butyl ether (MTBE) and 650 µL 25% methanol in water were added to extract the hydrophobic and hydrophilic compounds, respectively. 500 µL of the upper hydrophobic layer and 400 µL of the lower hydrophilic layer were transferred to separate autosampler vials and dried under nitrogen. The hydrophobic layer was reconstituted with 100 µL of methanol and the hydrophilic layer was reconstituted with 50 µL 5% acetonitrile in water. Both fractions were stored at −80 °C until LC/MS analysis.

#### Liquid Chromatography Mass Spectrometry

The hydrophobic fractions were analyzed using reverse phase chromatography on an Agilent Technologies (Santa Clara, CA) 1290 ultra-high precision liquid chromatography (UHPLC) system on an Agilent Zorbax Rapid Resolution HD SB-C18, 1.8um (2.1 x 100mm) analytical column as previously described (107, 108). The hydrophilic fractions were analyzed using hydrophilic interaction liquid chromatography (HILIC) on a 1290 UHPLC system using an Agilent InfinityLab Poroshell 120 HILIC-Z (2.1 x 100mm) analytical column with gradient conditions as previously described (109) with mass spectrometry modifications as follows: nebulizer pressure: 35psi, gas flow: 12L/min, sheath gas temperature: 275C, sheath gas flow: 12L/min, nozzle voltage: 250V, Fragmentor: 100V. The hydrophobic and hydrophilic fractions were run on Agilent Technologies (Santa Clara, CA) 6545 Quadrupole Time of Flight (QTOF) mass spectrometer. Both fractions were run in positive electrospray ionization (ESI) mode.

#### Mass Spectrometry Data Processing

Compound data was extracted using Agilent Technologies (Santa Clara, CA) MassHunter Profinder Version 10 software in combination with Agilent Technologies Mass Profiler Professional Version 14.9 (MPP) as described previously (47). Briefly, Batch Molecular Feature Extraction (BMFE) was used in Profinder to extract compound data from all samples and sample preparation blanks. The following BMFE parameters were used to group individual molecular features into compounds: charge state 1-2, with +H, +Na, +NH4 and/or +K charge carriers. To reduce the presence of missing values, a theoretical mass and retention time database was generated for compounds present in samples only from a compound exchange format (.cef) file. This .cef file was then used to re-mine the raw sample data in Profinder using Batch Targeted Feature Extraction.

An in-house database containing KEGG, METLIN, Lipid Maps, and HMDB spectral data was used to putatively annotate metabolites based on accurate mass (≤ 10 ppm), isotope ratios and isotopic distribution. This corresponds to a Metabolomics Standards Initiative metabolite identification level three (110). To improve compound identification, statistically significant compounds underwent tandem MS using 10, 20, and 40V. Fragmentation patterns of identified compounds were matched to either NIST14 and NIST17 MSMS libraries, or to the *in silico* libraries, MetFrag (111) and Lipid Annotator 1.0 (Agilent) (112).

#### Microbiome-associated metabolites

Microbiome-associated metabolites were defined using metabolites identified as significantly different in abundance between germ-free compared to humanized gnotobiotic mice and/or metabolites identified as microbially produced by the tool AMON (47).

For the gnotobiotic mouse analysis aqueous and lipid metabolites were analyzed separately. Metabolites that were present in <20% of samples were filtered out before analysis. Significant difference was determined using a Student’s t-test with FDR p-value correction. FDR-corrected p values < 0.05 were deemed significant. Significant metabolites also present in the human samples were retained for further analysis.

For the AMON-identified metabolites, the tool used an inferred metagenome, which was calculated using the PICRUSt2 QIIME2 plugin (49) and default parameters; a list of all identified KEGG IDs from the metabolite data (see metabolome methods); and KEGG flat files (downloaded 2019/06/10). AMON determined metabolites observed that could be produced by the given genes list. These metabolites were kept for analysis in addition to the gnotobiotic mouse identified metabolites. Those without any putative classification were removed from analysis.

### Microbiome Methods

#### Sample Collection, Extraction, and Sequencing

Stool samples were collected by the patient within 24 hours of their clinic visit on sterile swabs dipped into a full fecal sample deposited into a commode specimen collector. Samples were kept cold or frozen at −20°C during transport prior to being stored at −80°C. DNA was extracted using the standard DNeasy PowerSoil Kit protocol (Qiagen). Extracted DNA was PCR amplified with barcoded primers targeting the V4 region of 16S rRNA gene according to the Earth Microbiome Project 16S Illumina Amplicon protocol with the 515F:806R primer constructs (113). Control sterile swab samples that had undergone the same DNA extraction and PCR amplification procedures were also processed. Each PCR product was quantified using PicoGreen (Invitrogen), and equal amounts (ng) of DNA from each sample were pooled and cleaned using the UltraClean PCR Clean-Up Kit (MoBio). Sequences were generated on six runs on a MiSeq sequencing platform (Illumina, San Diego, CA).

#### Microbiome Sequence Processing and Analysis

Microbiome processing was performed using QIIME2 version 2018.8.0 (114). Data was sequenced across five sequencing runs. Each run was demultiplexed and denoised separately using the DADA2 q2 plugin (115). Individual runs were then merged together and 99% *de novo* OTUs were defined using vSEARCH (116). Features were classified using the skLearn classifier in QIIME2 with a classifier that was pre-trained on GreenGenes13_8 (117). The phylogenetic tree was building using the SEPP plugin (118). Features that did not classify at the phylum level or were classified as mitochondria or chloroplast were filtered from the analysis. Samples were rarefied at 19,986 reads. To reduce the number of comparisons within the microbiome, we binned highly co-correlating groups of measures within the data types into modules (Supplemental Table 2). These modules were defined using the tool, SCNIC (119). For statistical analysis features present in <20% of samples were filtered out.

### Bioinformatics

#### Module definition

Modules were called on microbiome and diet data. Modules were defined using the tool SCNIC (119). The q2-SCNIC plugin was used with default parameters for the microbiome data and standalone SCNIC was used for the diet data (https://github.com/shafferm/SCNIC). Specifically, for each data type SCNIC was used to first identify pairwise correlations between all features. Pearson correlation was used for diet and SparCC (120), which takes into account compositionality, was used for microbiome data. Modules were then selected with a shared minimum distance (SMD) algorithm. The SMD method defines modules by first applying complete linkage hierarchical clustering to correlation coefficients to make a tree of features. Modules are defined as subtrees where all pairwise correlations between all pairs of tips have an R value greater than defined minimum. The diet modules were defined using a Pearson r^2^ cutoff of 0.75. The microbiome modules were defined using a SparCC minimum r cutoff of 0.35. To summarize modules SCNIC uses a simple summation of count data from all features in a module. Application of SCNIC reduced the number of evaluated features from 6,913 to 6,818 for microbiome and 59 to 29 for diet data.

#### Statistical Analysis

All statistics were performed in R. For non-parametric tests Spearman rank correlation and Kruskal-Wallis test were used. For parametric tests linear models and Student’s t-test were used.

#### Data analysis tools

Metabolic disease score was calculated using PCA in R with prcomp. Data was scaled using default method within the prcomp library. All random forest analysis tools were used in R. Standard random forest was performed using the randomForest function. Variable selection was performed in R using the tool VSURF (38). Interaction analysis was performed in R using the tool iRF (42).

### Data Availability

Microbiome data is available at EBI/ENA (https://www.ebi.ac.uk/ena) accession number: ERP125300. Immune and diet data are available along with the microbiome data as associated metadata. Metabolomics data will be available on Metabolomic Workbench (https://www.metabolomicsworkbench.org) upon publishing. Until publicly available it is available upon request.

## Acknowledgements

We would like to thank our study participants for contributing their samples and time to this study. We would also like to thank Christine Griesmer for her contributions to subject recruitment and Brandi Wagner and Maggie Stanislawski for insights into statistical analysis methods.

## Author contributions

AJSA analyzed and interpreted all data. AJSA and CAL wrote the manuscript. NR guided generation and interpretation of metabolomics data. KQ and KAD prepared, ran, and processed metabolomics. KQ ran metabolic pathway analysis. SXL prepared and conducted mouse experiments. JMS ran immunological assays. NMN prepared and ran sequencing and coordinated fecal sample and metadata collection from study subjects. SF recruited subjects, collected samples, and maintained regulatory compliance. TJM and JH collected and aided in interpretation and processing of diet data. CAL, BEP, and TC conceptualized and led the study.TC guided all clinical data collection and subject recruitment and provided clinical insight into study populations. BEP guided generation and interpretation of immune data. CAL guided microbiome data generation and multi’omic data analysis. All authors read and approved the final manuscript.

## Supplemental Tables

**Supplemental Table 1.** Pairwise test p-values for Table 1 demographics

**Supplemental Table 2.** Study measures by datatype and SCNIC-calculated modules for microbiome and diet survey data

**Supplemental Table 3.** VSURF-selected features: correlation with metabolic disease score, edge table for inter-variable correlations, and iRF analysis results

**Supplemental Table 4.** Gnotobiotic mouse experiment set-up and western diet

**Supplemental Table 5.** Microbiome-associated metabolites list with source of identification, metabolites correlating with metabolic disease score and mBrole pathway analysis results

## Supplemental Figures

**Supplemental Figure 1. Metabolic disease score cutoff calculations using regressions between metabolic disease score and the metrics used in the PCA analysis.** Triglycerides, fasting glucose, HDL and LDL have well-define clinical cut-offs for high values and were used to calculate the healthy-unhealth cutoff. Linear model was calculated modeling metabolic disease score by each marker. The high value intercept of the regression line is marked with a dotted line and value annotated on the plot. The solid line is the defined healthy-unhealthy cutoff calculated as the mean of the four cutoff values. P values are from the linear model.

**Supplemental Figure 2. Histogram of percent variation explained in permuted VSURF.** Metabolic disease score was permuted 1,000 times and passed through VSURF. The resulting variables were run through a standard random forest and the percent variation explained was calculated. The blue line represents the percent variation explained for the true VSURF. P value was calculated using a one tailed test.

**Supplemental Figure 3. Correlation network of VSURF-selected variables.** Correlation network of **A)** all VSURF-selected variables, **B)** neighboring nodes of dietary fiber, and **C)** neighboring nodes of age. All Spearman rank correlations with an FDR p < 0.25 are shown. See Supplemental Table 3 for the edge table and Supplemental Table 3 for the node table.

**Supplemental Figure 4. Metabolic disease associated metabolites by model type with age accounting.** Venn diagram summarizes all significant results (FDR-corrected p-value < 0.05) of each test and the metabolites that overlap between tests that do and do not account for age correction.

