## Supplementary figures and images for "Systems analysis of gut microbiome influence on metabolic disease in HIV and high-risk populations"

### Supplemental Figure 1

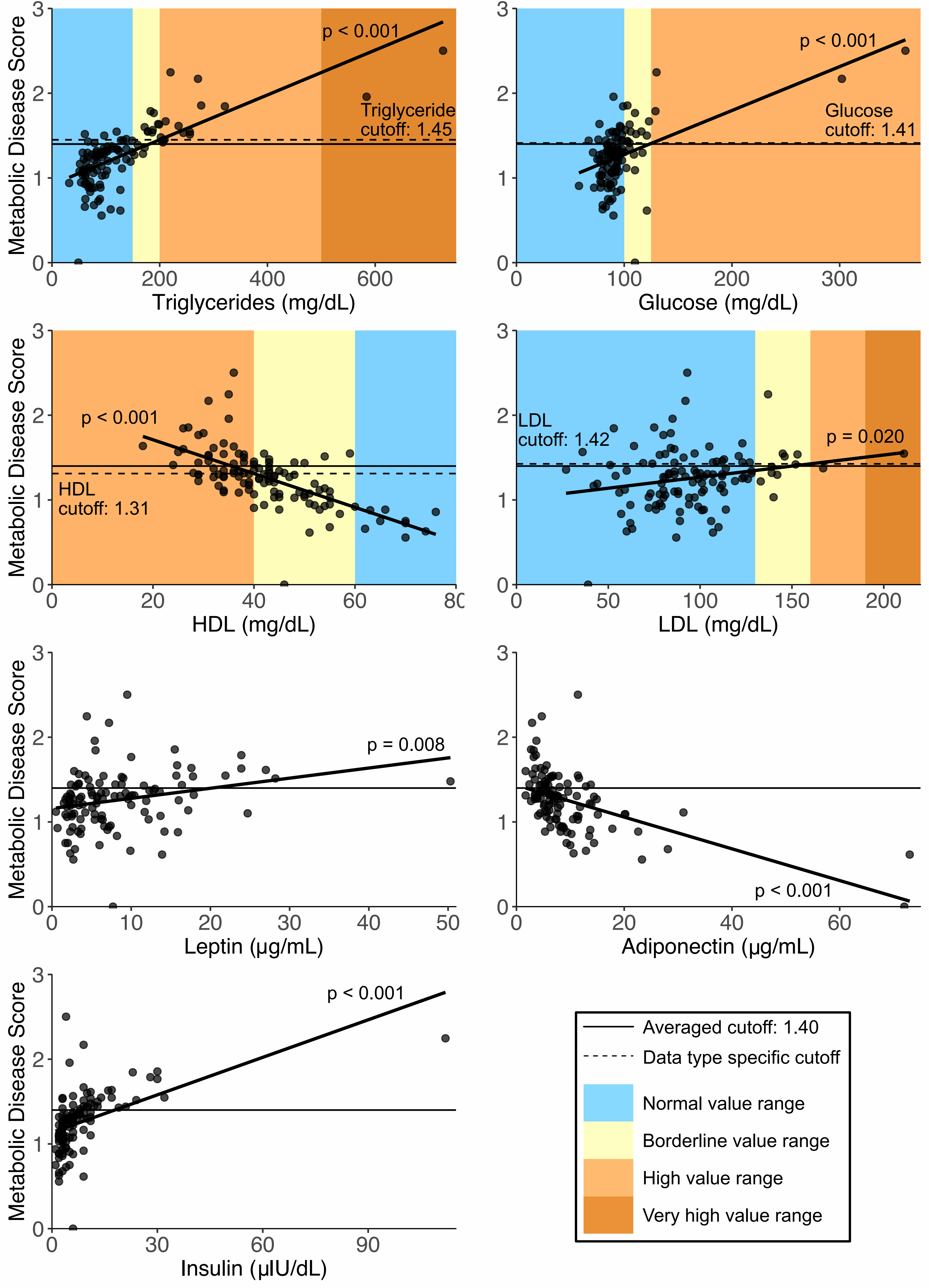

### Supplemental Figure 2

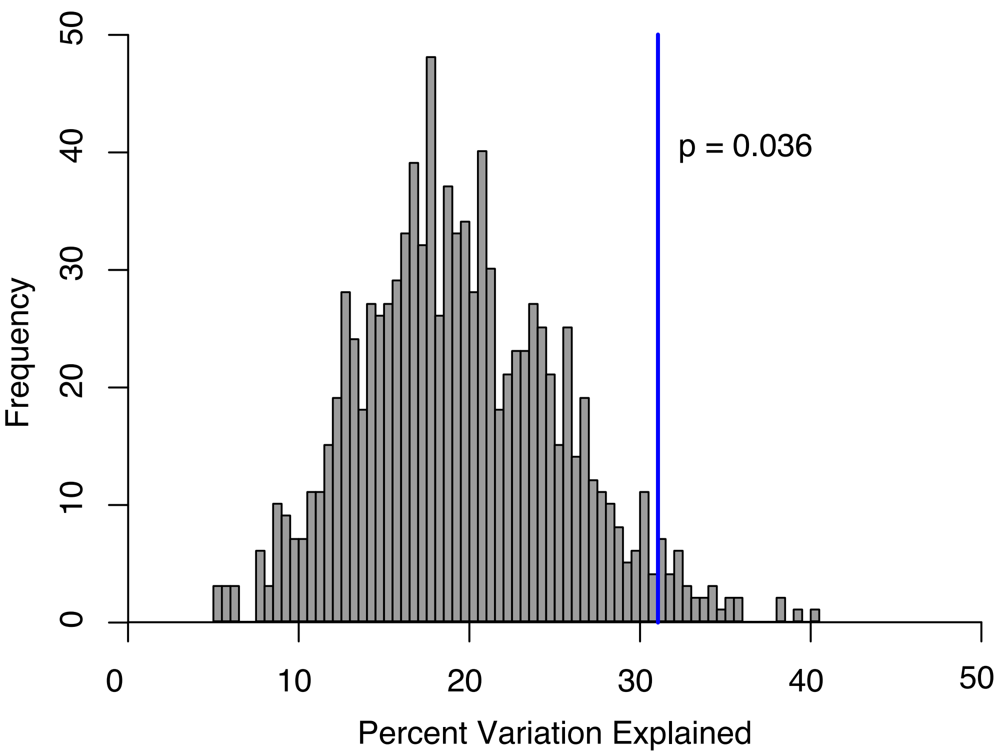

### Supplemental Figure 3

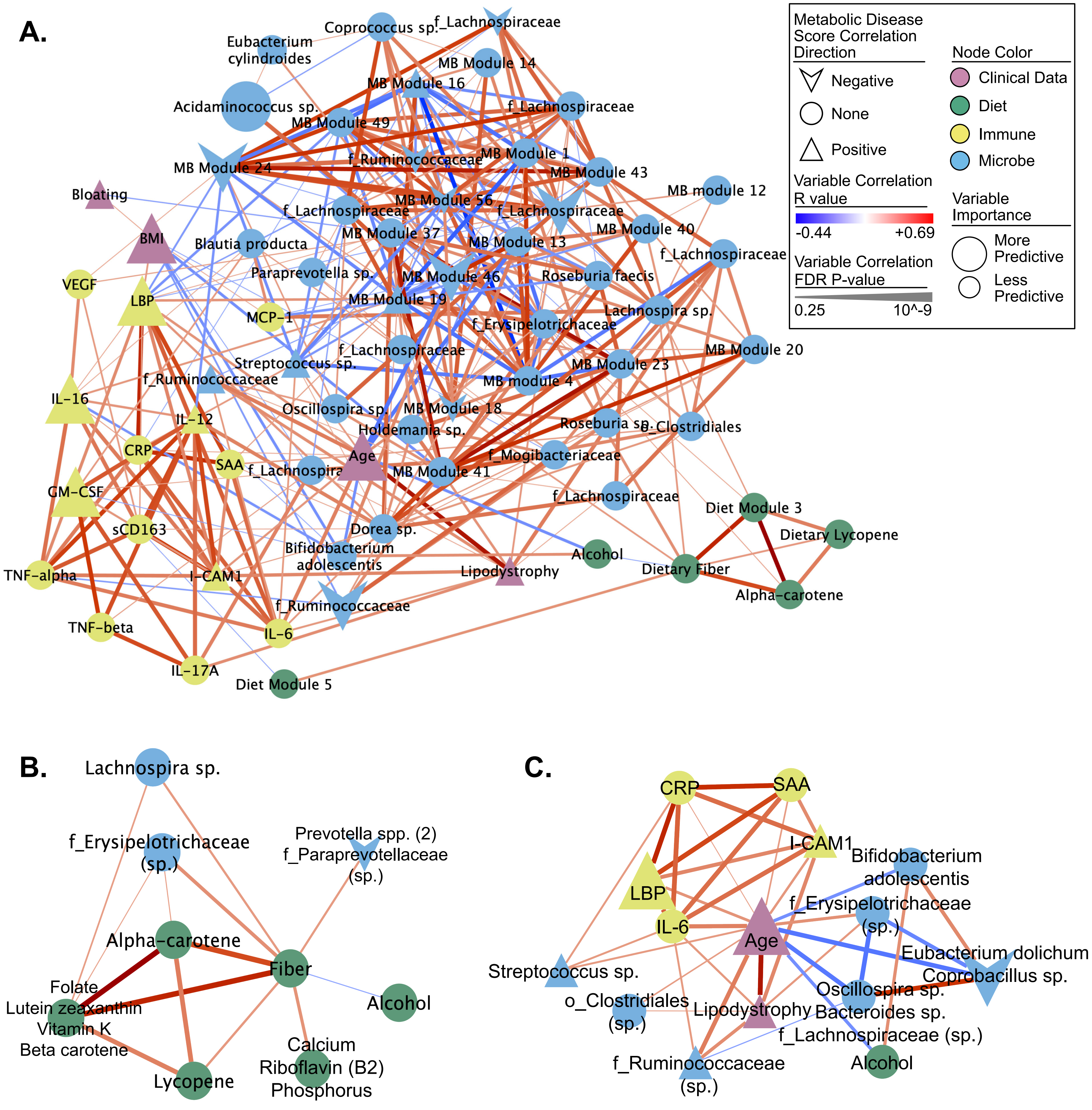

### Supplemental Figure 4

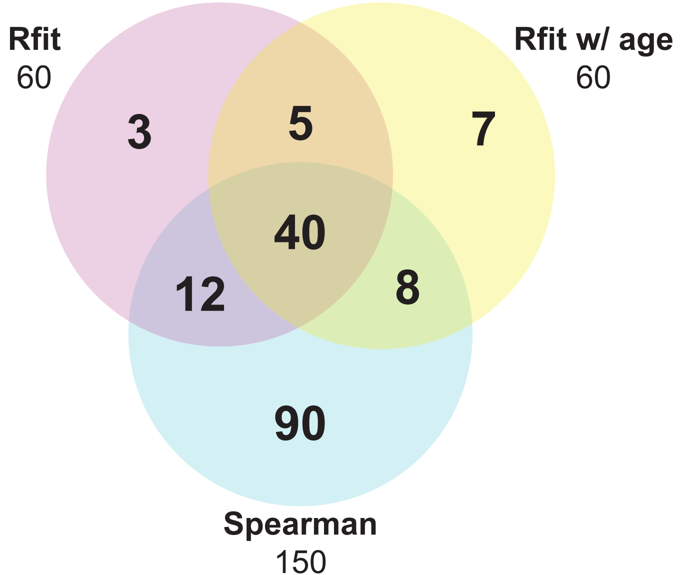
